# Smooth pursuit eye movements contribute to long-latency reflex modulation in the lower extremity

**DOI:** 10.1101/2025.01.12.632642

**Authors:** Oindrila Sinha, Angus Peter Muttee, Jia-Hao Wu, Matteo Bertucco, Isaac Kurtzer, Tarkeshwar Singh

## Abstract

Somatosensory mediated reactions play a fundamental role in adapting to environmental changes, particularly through long-latency responses (LLRs)—rapid corrective muscle responses (50-100ms) following limb perturbations that account for limb biomechanics and task goals. We investigated how smooth pursuit eye movements (SPEM), which are slow eye movements used to track moving objects, influence LLRs of the upper and lower limb during mechanical interactions with moving objects. In the first experiment, participants stood and stabilized their arm against a colliding virtual object. This occurred while subjects either visually pursued the moving object or fixated a central location. The robot occasionally applied a mechanical perturbation to the arm either 200ms or 60ms before the anticipated collision. As in previous studies, LLRs were observed in leg muscles to a perturbation of the upper limb. Moreover, leg LLRs were modulated by gaze, being larger during pursuit than fixation but only during the late perturbations. This timing-specific modulation aligns with previous reports of policy transitions in feedback control roughly 60ms before impact. Upper limb LLRs were not significantly impacted by gaze. This lack of modulation could reflect the context of upright stance, so we conducted a second experiment which was the same in all ways except that the subjects remained seated. Again, the upper limb LLRs were not impacted by gaze. The selective impact of gaze modulation on stance control highlights the sophisticated nature of coordinating eye movements, arm control, and whole-body postural responses.

**NEW and NOTEWORTHY:** Smooth pursuit eye movements are used to track moving objects. We show for the first time that smooth pursuit eye movements contribute to modulation of long-latency reflexes in the lower limb during virtual object interactions in upright stance. These findings suggest a neurophysiological link between predictive control of eye movements and feedback control of upright stance.

## INTRODUCTION

Our sensorimotor system is well adapted to the complex and changing environment we inhabit. This competence is exemplified by a waiter receiving an unexpected bump and stabilizing their tray based on its content, the physical surroundings, and even shifting their weight to remain upright. Experiments show that such sophistication is within the long-latency reflex (LLR), an epoch occurring ∼50-120 ms post-perturbation. LLRs can be elicited in muscles remote to the local stretch, such as lower extremity LLRs with upper limb perturbations (1, 2). LLRs are also modulated by task goals and experimental contexts (3, 4), and are highly influenced by visual information (5–7) reflecting the funneling of information from frontoparietal cortical areas onto motor cortical circuits which generate the LLR.

The modulatory impact of vision on somatosensory evoked LLRs includes scaling to continuous motion signals, such as the direction and coherence of random-dot kinematograms as used in studies of decision-making studies (7). The key cortical substrate for motion processing is middle temporal complex (MT+) and its impact on LLRs is especially notable since MT+ plays a crucial role in smooth pursuit eye movements (SPEM) used to track moving objects (8, 9). During SPEM, an efference copy of the extraretinal signal associated with eye movements has been hypothesized to contribute to motion perception (11–13), to guide upper limbs actions such as interceptive hand movements (14), and anticipatory posture stabilization during interactions with moving objects (15, 16).

The broad influence of SPEM on limb motor control and the linkage of MT+ to LLR modulation suggests an important but untested possibility: LLRs may be modulated by gaze behavior. To test this hypothesis, we examined healthy individuals as they stood upright (Exp.1) and countered a virtual object that collided with their arm, and either freely pursued the moving object with their eyes or fixated a central location. On a random subset of trials, we applied mechanical perturbations to the arm to evoke LLRs of the upper and lower limbs. We applied mechanical perturbations either ∼200 ms or ∼60 ms prior to the impending collision as previous work on the role of vision for ball catching showed that stretch reflexes are modulated ∼60 ms before collision to increase limb impedance (17, 18). We found that the leg LLRs, but not arm LLRs, were modulated by gaze behavior immediately before the impeding collision. The lack of gaze modulation for the arm could reflect the context of upright stance. Arm and leg reflexes to the same perturbation can be highly modified between many postural contexts such as seated versus free-standing and free-standing versus standing with a trunk support or holding a stable bar (19). We therefore performed an additional identical experiment with participants in a seated position (Exp. 2). Consistent with our initial findings, arm LLRs showed no modulation by gaze behavior.

## METHODS

### Participants

Twenty-nine healthy right-handed participants completed the study: seventeen in Experiment 1 (age: 23.3 ± 1.2 years; 9M/8F) and twelve in Experiment 2 (age: 22.5 ± 1.05 years; 6M/6F). Participants provided written informed consent and were compensated ($10/h). The study was approved by Penn State’s Institutional Review Board.

### Apparatus

Participants interacted with a Kinarm Endpoint robot (Kinarm, Canada) integrated with an SR EyeLink 1000 Remote eye-tracker (SR Research, Canada). Visual stimuli were projected (120 Hz) onto a semi-transparent mirror via a VPixx display (VPixx Technologies, Canada). Direct vision of the hand was occluded (Fig. 1A). In Exp. 1, participants stood upright on a force plate (Bertec, USA), head tilted ∼30° forward, while manipulating the robot handle. EMG data were recorded wirelessly (Trigno, Delsys Inc, USA) from tibialis anterior muscles and the right biceps and triceps brachii. In Exp. 2, participants were seated, and EMG was recorded only from upper extremity muscles. EMG data and center of pressure were sampled at 1,000 Hz. Hand forces were measured (1,000 Hz) using an ATI-Mini40 force transducer (ATI Automation, USA). The monocular eye-tracker, mounted 80 cm from participants, recorded left eye movements at 500 Hz (accuracy: 0.25-0.5°, data loss: 1.6 ± 0.9%, (20)) and was calibrated using Kinarm’s calibration routine. Experiments were conducted in dim light. Participants held only the robot handle and kept their forehead slightly away from the head support.

**Figure 1:**
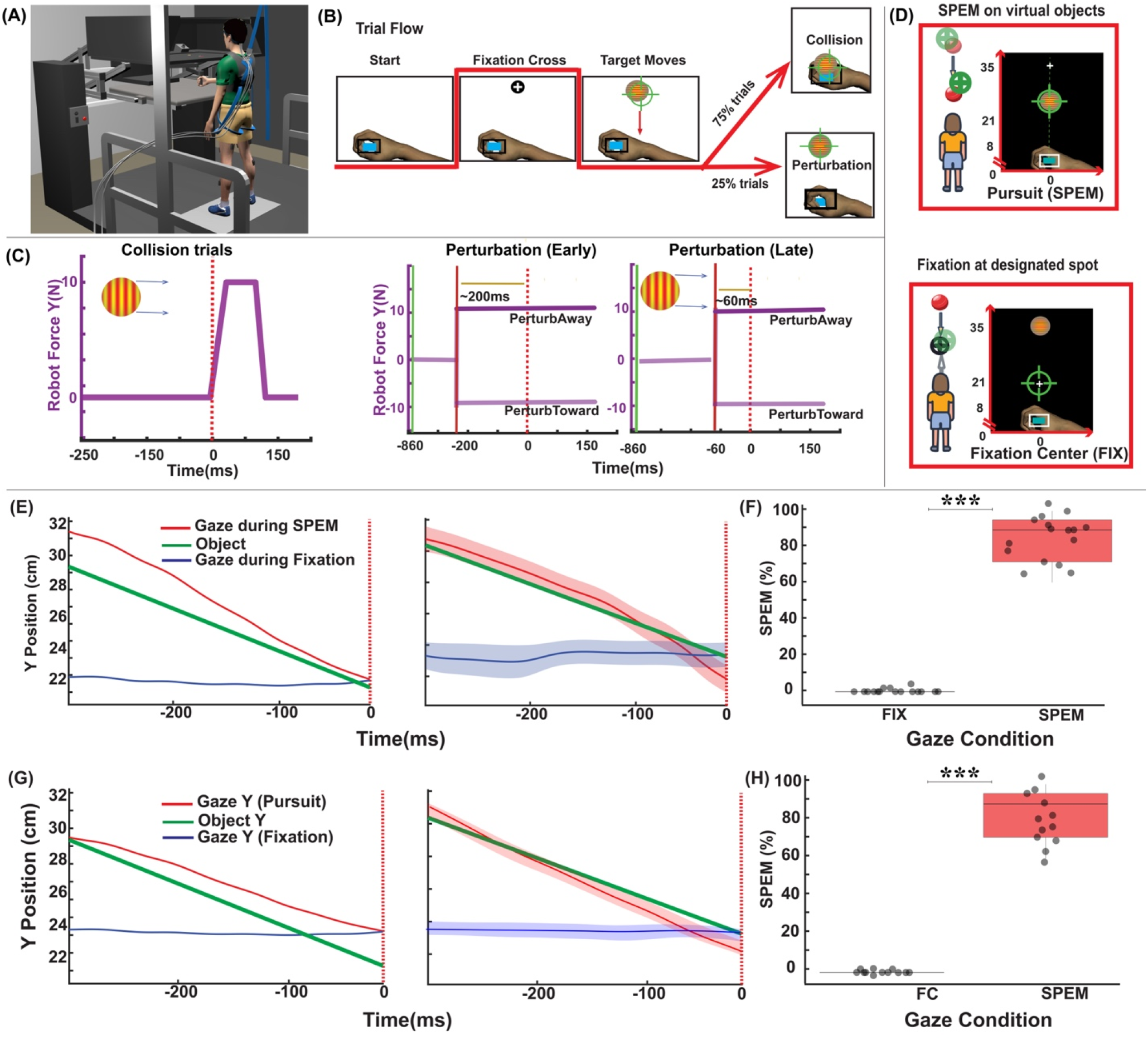
Experimental setup, protocols, and gaze data. (A) Participants stood at a KINARM endpoint robot with force plate. (B) Trials began with hand positioning in a virtual target, followed by fixation cross and moving object presentation (25 cm/s). (C) Force profiles: collision trials (75%) delivered 90 ms trapezoidal forces; perturbation trials (25%) applied step forces either early (Early, ∼200 ms pre-contact) or late (Late, ∼60 ms pre-contact) in anterior-posterior directions. (D) Participants completed 10 randomized blocks each of smooth pursuit (SPEM) and central fixation (FIX) conditions. (E) and (G) Single trial example (left panel) and mean gaze position (pursuit: red, fixation: blue) and object position (green) along the y-axis for all participants (right panel) in Exps. 1 & 2, respectively. (F) and (H) Pursuit percentage during fixation (FIX) and pursuit (SPEM) conditions show that participants tracked moving objects with SPEM during pursuit condition in Exps. 1 & 2, respectively.

### Experimental task

Participants maintained their hand position within a virtual target (1×1 cm rectangle) while standing (Exp. 1, Fig. 1A) and sitting (Exp. 2). Each trial began with cursor positioning at a start location, followed by fixation on a cross (Fig. 1B). After the cross turned off (250-750 ms random delay), a virtual object appeared 32 cm away and moved toward the participant’s hand position at 25 cm/s. In 75% of the trials, when the object collided with the hand, the robot applied a 10 N force impulse in the posterior direction (10 ms rise and fall times, 90 ms plateau) to simulate a collision (see Fig. 1C). Participants were instructed to maintain hand position within the target while resisting the force impulse during collision. In the remaining 25% of trials, mechanical perturbations (10N, 10 ms rise time, 90 ms plateau) were applied before anticipated object collision, either 200 ms (Early) or 60 ms (Late) ms before contact, either towards (PerturbToward) or away (PerturbAway) from the body. Perturbations were delivered in the anterior or posterior direction. The order of the trials was randomized within a block.

### Experimental conditions

We used two gaze conditions in each experiment (central fixation, FIX; and smooth pursuit eye movements, SPEM). In the smooth pursuit (SPEM) condition, participants were instructed to pursue the object all the way till contact. In the Fixation (FIX) condition, the visual fixation point was positioned 5 cm anterior to the start location along the body midline (see Fig. 1D).

Exp. 1 comprised 10 blocks of 32 trials per gaze condition (640 total trials), with 25% (160 trials) involving perturbations. These were equally divided between ‘PerturbToward’ and ‘PerturbAway’ (80 trials each). Analysis focused exclusively on ‘PerturbToward’ trials (40 ‘Early’ and 40 ‘Late’), comparing responses between central fixation (FIX) and pursuit (SPEM) conditions. ‘PerturbAway’ trials were excluded due to poor signal-to-noise ratio in the gastrocnemius and biceps muscles. Exp. 2 used a modified protocol with 28 trials per block, implementing only ‘PerturbToward’ perturbations.

### Study design

Participants began the study with 16 EMG normalization trials. Here they encountered a 10N static background force directed either towards or away (8 trials each) and maintained their arm position for 2.5 seconds. They then performed 10 blocks of FIX and 10 blocks of SPEM gaze conditions, randomly ordered across participants.

### Data recording and analyses

EMG signals were band-pass filtered (20-450 Hz), rectified, and smoothed using a 10-point moving average (for plotting). After baseline subtraction, signals were normalized to root mean square (RMS) values from normalization trials. For each gaze and experimental condition, trials were ensemble-averaged before quantifying response intervals. The windows for the upper and lower limb LLRs was 50-100 ms and 75-120 ms, respectively, to account for conduction delays (2). The voluntary response windows were 100-150 ms for the upper extremity and 120-150 ms for the lower extremity. The mean EMG within these windows was compared across gaze conditions.

Gaze data were pre-processed following established protocols (21, 22). Point-of-regard (POR) data were low-pass filtered (15 Hz), converted to spherical coordinates, and transformed to ocular kinematics. Angular gaze speed was computed using a Savitzky-Golay filter (6th-order, 27-frame window). Gaze events (saccades, fixations, smooth pursuits) were classified using a threshold-based algorithm. Figure 1E shows gaze and target position along the Y-axis for two exemplary trials in the left panel and the mean across participants in the right panel. Kinetic and kinematic data were low-pass filtered at 50 Hz and 15 Hz, respectively, using a double-pass, zero-lag, third order Butterworth filter. Center of pressure (COP) in the x (mediolateral)- and y (anterior/posterior)-directions was calculated with the following equations:

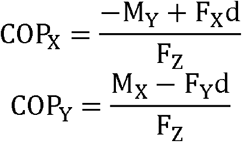

Where F_x_, F_y_, and F_z_ represent ground reaction forces in the x-, y-, and z-directions, respectively, and M_x_ and M_y_ denote moments about the x- and y-axes. ‘d’ signifies the distance from the top of the force plate to its center.

### Statistics

Data were tested for normality, homoscedasticity, and sphericity. If any assumption was not met, data were log-transformed. A repeated measures ANOVA examined effects of Gaze Condition (FIX, SPEM) and Perturbation Timing (Early, Late) on EMG amplitudes, center-of-pressure excursions, and hand force signals, with participants as random effects. Effect sizes were quantified using generalized η^2^. Post-hoc comparisons were done using Holm-corrected pairwise tests (α = 0.05).

## RESULTS

To test our hypothesis, we constrained the gaze in the Fixation (FIX) condition and allowed participants to freely pursue the moving object in the Pursuit (SPEM) conditions. Eye-tracking data confirmed successful gaze manipulation between conditions. In Exp. 1, during Fixation (FIX), smooth pursuit duration averaged 0.0 ± 0.1%, compared to 87.5 ± 1.3% during Pursuit (SPEM) blocks (Fig. 1F). In Exp. 2, smooth pursuit duration averaged 0.0 ± 0.4% during FIX (Figs. 1G and 1H), compared to 87.1 ± 1.2% during Pursuit (SPEM) blocks.

Pre-perturbation EMG amplitudes were comparable between gaze conditions across all recorded muscles. This baseline similarity was critical for comparing feedback responses, as short-latency response amplitudes typically scale with pre-perturbation activation, unlike long-latency responses (23, 24). Although short-latency responses were not our primary focus, controlled baseline activity enabled valid comparison of feedback responses. The pre-perturbation and pre-collision anticipatory EMG activity is shown in the supplementary material (Figs S1 and S2).

### Constraining eye movements did not affect LLR amplitudes in the triceps brachii

In Exp. 1, long-latency feedback responses in triceps brachii showed no significant gaze-dependent modulation during either the early or the late perturbations [Early: F(1,16) = 2.3, p = 0.7, η^2^ = 0.01; Late: F(1,16) = 6.1, p = 0.4, η^2^ = 0.08]. Similarly, in Exp. 2, long-latency feedback responses in triceps brachii showed no significant gaze-dependent modulation during either the early or the late perturbations [Early: F(1,11) = 0.2, p = 0.6, η^2^ = 0.02; Late: F(1,11) = 0.5, p = 0.9, η^2^ = 0.02]. This suggests that extraretinal signals from smooth pursuit eye movements did not influence upper limb LLR responses (Fig. 2A-C).

**Figure 2:**
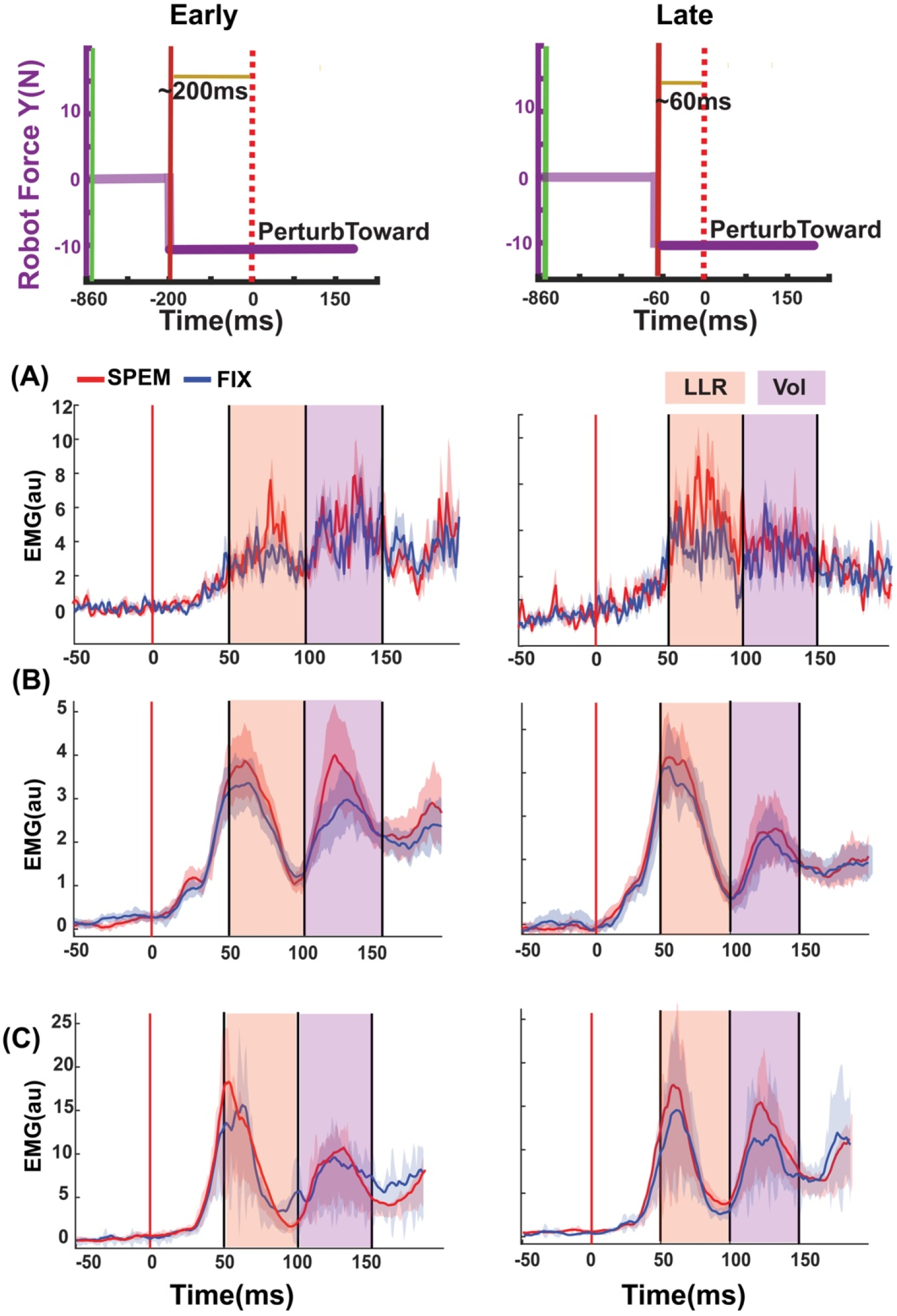
Long-latency reflexes in triceps brachii. The top panel shows the direction and timing of the perturbation applied by the robot. Left plot shows early (Early) and right plot shows late (Late) perturbations. Representative single-participant (A) and group-averaged (B) data showing long-latency reflexes (LLR, orange) and voluntary responses (purple) windows for Exp. 1. C) Group-averaged data for Exp 2.

### Constraining eye movements significantly decreased the amplitude of LLR in the lower limb muscles

LLR responses in bilateral tibialis anterior muscles showed gaze-dependent modulation, particularly during perturbations soon before the anticipated contact (Late perturbation). Figure 3A-B illustrates a representative participant’s data (upper panels) and the group average (lower panels) for left and right tibialis anterior. The center of pressure adjustments (COP_y_) remained unaffected (Supplementary Materials, Figure S3), suggesting selective influence of extraretinal signals on long-latency feedback responses, but not on overall metrics of posture stabilization.

**Figure 3:**
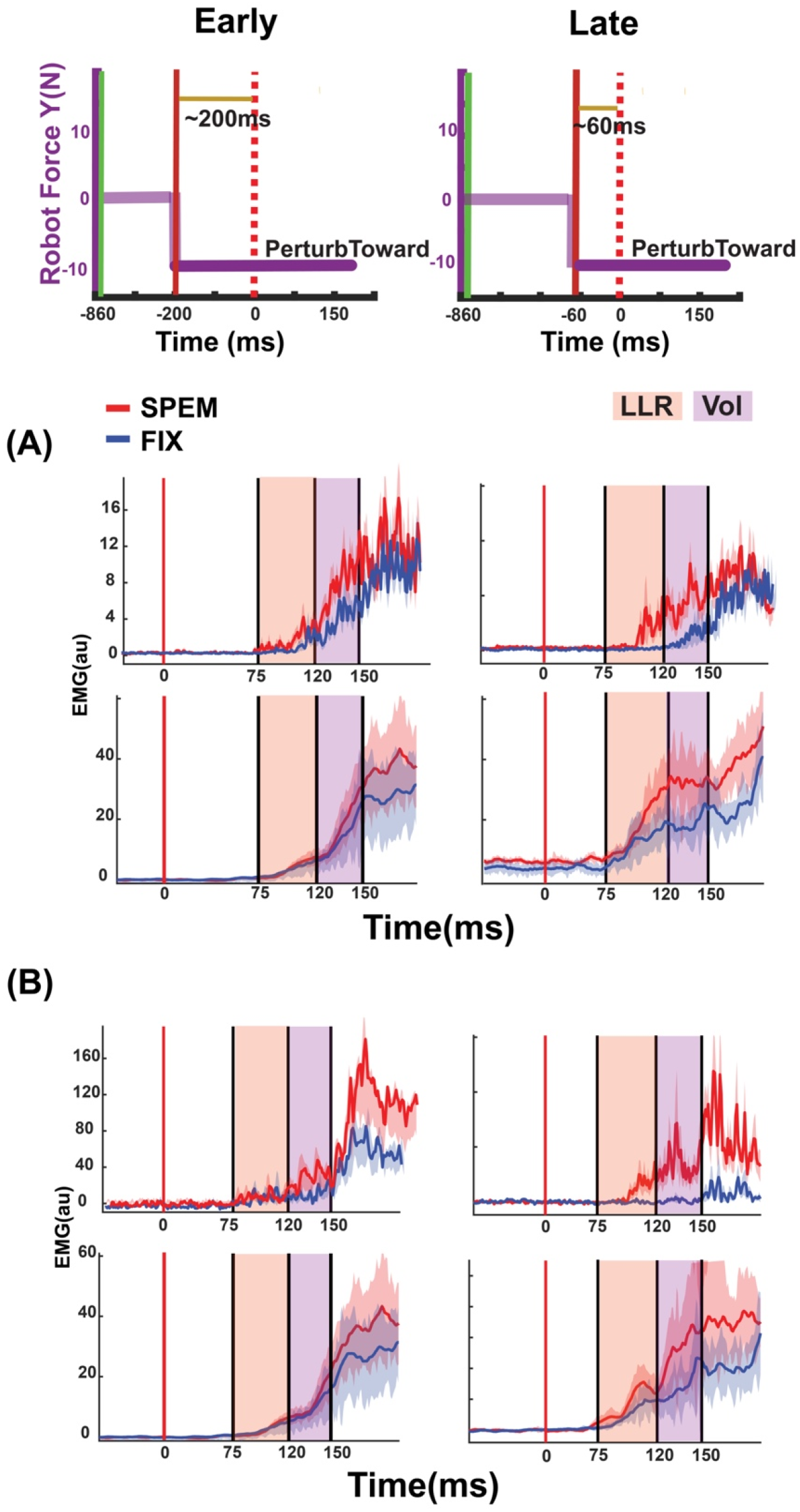
Gaze-dependent modulation of lower limb reflexes in Exp. 1. The top panel shows the direction and timing of the perturbation applied by the robot. EMG activity during early (Early, left) and late (Late, right) perturbations. (A) Left tibialis anterior: Single-participant example (top) and group average (n=17, bottom) showing gaze-dependent modulation in long-latency (orange) and voluntary (purple) windows. (B) Right tibialis anterior: Similar modulation pattern in single-participant (top) and group data (bottom).

Mean long-latency reflex responses in tibialis anterior muscles were log-transformed and compared across the two gaze conditions at both perturbation times. Right tibialis anterior showed significantly larger long-latency feedback responses during pursuit compared to fixation [F(1,16) = 20, P =0.03, η^2^ = 0.2; Fig. 4B]. During late perturbations, the differences between the gaze conditions in the left tibialis muscles approached significance [F(1,16) = 8.2, P = 0.1, η^2^ = 0.03; Fig. 4A]. These results indicate timing-specific modulation of postural reflexes by extraretinal signals, particularly in the right leg.

**Figure 4:**
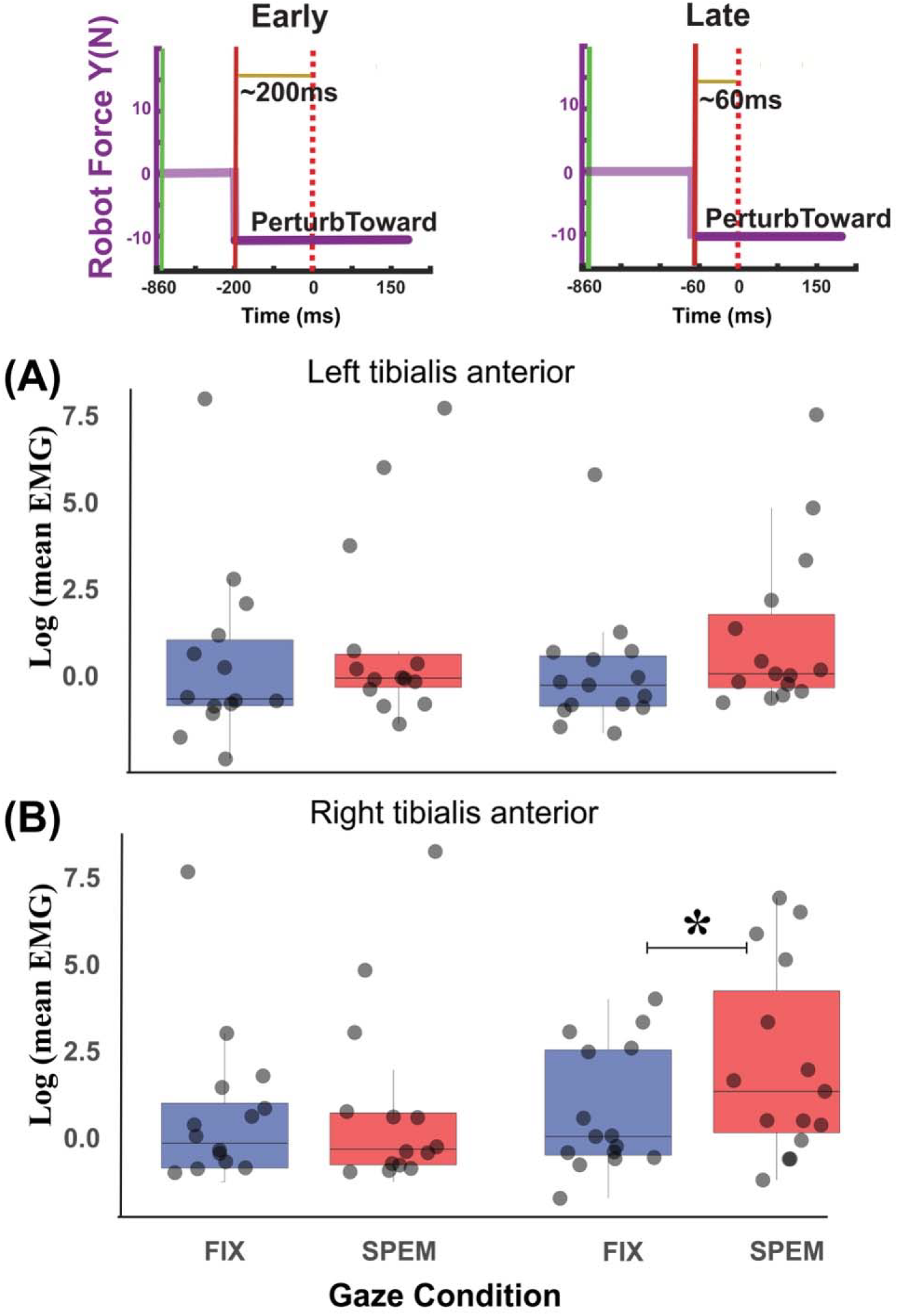
Long-latency reflex modulation in Exp. 1. Box plots showing LLR amplitude (log-transformed EMG, 75-120 ms post-perturbation) during early (EARLY, left plots) and late perturbations (Late, right plots) for (A) left and (B) right tibialis anterior muscles. Gaze conditions: pursuit (SPEM) vs. fixation (FIX). *P < 0.05.

## DISCUSSION

Our findings replicated and extended multiple research domains. First, we demonstrated that upper limb perturbations elicited long-latency reflexes (LLRs) in the lower limb (Exp. 1), consistent with previous observations in postural and limb perturbation studies (2, 25) and the broader framework of multi-segment integration in rapid feedback control (3). Second, we identified gaze-dependent modulation of leg LLRs, with enhanced responses during tracking of approaching objects with smooth pursuit eye movements (SPEM) compared to central target fixation. This gaze dependency aligns with extensive research on SPEM’s role in motion processing (26) and interception movements (14), where SPEM demonstrates performance advantages and is preferentially selected in free-choice paradigms (15).

Our previous work established that smooth pursuit eye movements (SPEM) lead to different patterns of anticipatory upper limb control when a seated subject must exert a matching force pulse against a colliding virtual object. Specifically, SPEM amplifies the anticipatory force magnitude compared to central fixation (15). Notably, peripheral fixation yields response patterns similar to SPEM, reflecting enhanced motion sensitivity in peripheral vision (16). The observed differential effects of SPEM versus central fixation on reactive compensation to mechanical perturbations complement our earlier findings on anticipatory compensation, suggesting substantial mechanistic overlap and supporting broader theoretical frameworks (27).

The gaze-dependency of leg LLRs manifested during the period closest to anticipated collision, but not the earlier epoch. This temporal specificity indicates that the effect is not attributable to general SPEM-related upregulation, which was constant throughout, nor to global visual differences previously documented in vision versus no-vision LLR paradigms, where responses were enhanced in the no-vision condition (28). Instead, the observed modulation was epoch-specific, aligning with evidence for temporal adjustments in somatosensory processing (29), reflex engagement (5), and control strategies immediately preceding mechanical interactions. The persistence of this pattern under visual occlusion remains unexplored and is an obvious next step.

Our findings likely reflect the integrative function of the middle temporal complex (MT+) in processing both retinal and extraretinal motion signals during smooth pursuit eye movements (31). MT+ engagement has been demonstrated in both small-object tracking and large-field motion processing tasks, such as random dot kinematograms (32), with its causal role in motion processing evidenced by post-ablation performance deficits (33). Building on Selen et al.’s (7) work that long-latency feedback response gains scale with decision time, we propose that MT+ motion signals may modulate premotor regions either directly or via parietal areas associated with perceptual evidence accumulation (34).

Our results strengthen evidence for MT+’s modulatory role in LLR gain control through integrated extraretinal and retinal signal processing and subsequent PMd signaling. However, our experimental design’s temporal resolution, limited to two discrete perturbation timepoints, precluded investigation of continuous MT+ and premotor dynamics. Additionally, our task demanded discrete sensorimotor responses unlike random-dot motion paradigms that require ongoing evidence accumulation. Despite these methodological distinctions, both paradigms implicate MT+ in LLR gain modulation, suggesting a conserved role in sensorimotor integration. It should be noted that our conclusions regarding MT+’s specific role in this process remain speculative without direct neural recordings and would benefit from further investigation using complementary methodologies.

The selective modulation of lower extremity feedback responses (Figs 3 & 4) was unexpected but aligns with established literature demonstrating context-dependent differential control of upper and lower limbs, such as during fixed-support versus surface perturbation conditions (35). This limb-specific modulation suggests preferential gaze-dependent postural control in the lower extremities during standing.

We hypothesized that gaze-dependent modulation would similarly affect upper limb long-latency reflexes (LLRs) during seated reaching tasks. However, Experiment 2 revealed no significant gaze-dependent effects on upper limb LLRs. Given that LLRs demonstrate task-dependent modulation and contribute to limb stabilization, gaze-dependent modulation may emerge under more challenging conditions requiring dynamic postural control and impact anticipation. Future experiments will investigate whether upper limb LLRs exhibit gaze-dependent modulation during interceptive actions that demand active postural stabilization against external perturbations.

Taken together, the gaze-dependent modulation of LLRs was restricted to lower extremity muscles, suggesting a hierarchical organization of postural control. This selective modulation may reflect the prioritization of postural stability (36), where SPEM-associated extraretinal signals preferentially influence postural control circuits. The anatomical basis for this modulation could involve known projections from MT+ or superior colliculus—key nodes in the SPEM network—to the pontine nuclei and reticular formation, respectively. The precise neural mechanisms underlying this selective modulation and the relative contribution of these pathways to postural stability require further investigation.

## Supporting information

https://doi.org/10.6084/m9.figshare.28548989.v1

## ACKNOWLEDGEMENTS

We thank Ana Spasic for help with data collection.

## SUPPLEMENTAL MATERIAL

Supplemental Figs. S1-S3: https://doi.org/10.6084/m9.figshare.28548989.v1

## DATA AVAILABILITY

All the data and analyses codes from the project are publicly available at DOI: 10.17605/OSF.IO/S8W2Y (https://osf.io/s8w2y/).

