## Supplementary material for "Smooth pursuit eye movements contribute to long-latency reflex modulation in the lower extremity": https://doi.org/10.6084/m9.figshare.28548989.v1

**Gaze condition did not significantly affect EMG activity, hand forces, movement patterns, or center of pressure during either collision or perturbation trials**

In the collision trials (Fig. S1), analysis of anticipatory activity (500 ms pre-collision) revealed no significant gaze-dependent effects across multiple measures. ANOVA showed no effect of gaze condition on triceps EMG [F(1,33) = 0.03, p = 0.88, η² = 0.07; Figure S1A], hand force [F(1,33) = 0.08, p = 0.65, η² = 0.08; Figure S1B], hand position [F(1,33) = 0.02, p = 0.98, η² = 0.05; Figure S1C], bilateral tibialis anterior EMG [left: F(1,33) = 0.01, p = 0.49, η² = 0.1; right: F(1,33) = 0.3, p = 0.56, η² = 0.2; Figures S1D,E], or center of pressure [x: F(1,33) = 10, p = 0.71, η² = 0.1; y: F(1,33) = 11, p = 0.89, η² = 0.2; Figure S1F].

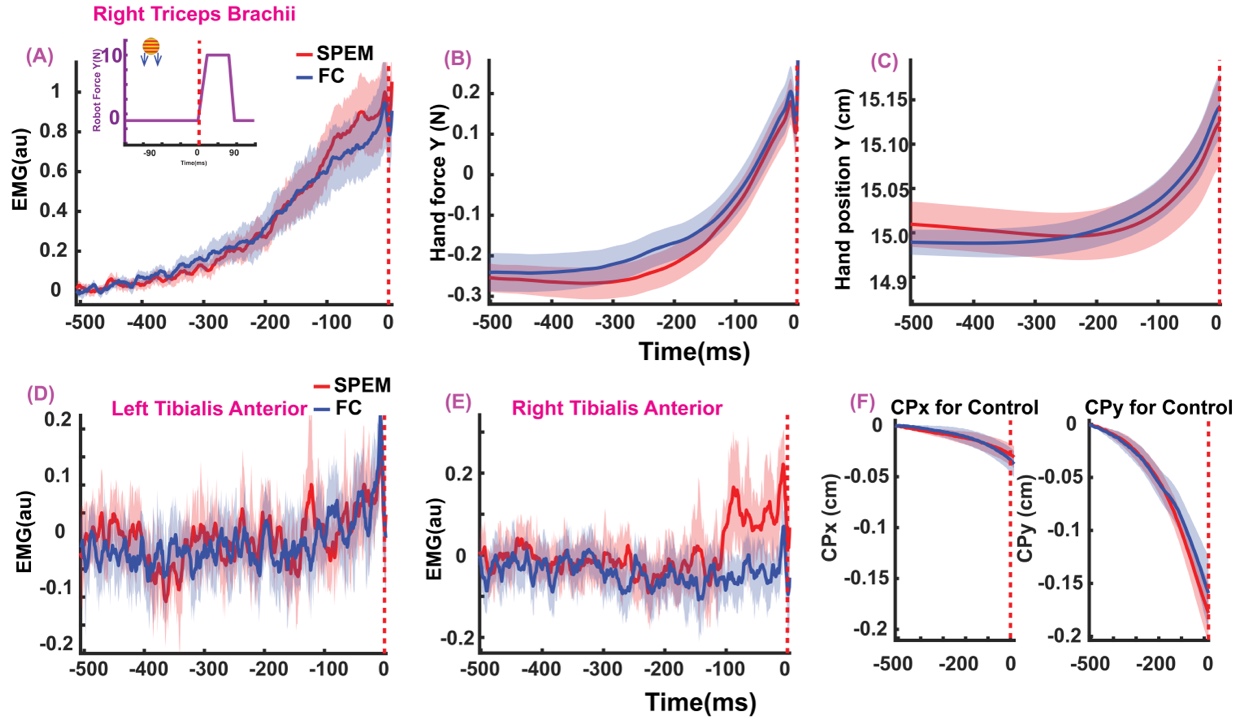

Figure S1: Pre-collision anticipatory activity. (A) Triceps EMG shows similar pre-collision increase for both gaze conditions (pursuit: red, fixation: blue). Hand force (B) and hand position (C) in Y-direction display minimal pre-collision increase across both gaze conditions. (D, E) Bilateral tibialis anterior EMG shows minimal pre-collision changes. (F) Center of pressure along the X-direction did not change much, but along the Y-direction drifted about a mm towards the participant prior to the collision in both gaze conditions.

Data in Figure S1 show increased pre-perturbation baseline activity in the triceps muscle, producing minimal hand movement toward the target (~1.5 mm), suggesting upper extremity co-contraction prior to collision. However, poor signal-to-noise ratio in in the antagonist biceps EMG prevented confirmation of co-contraction.

In the remaining 25% of the perturbation trials, ANOVA revealed no gaze-dependent effects on pre-perturbation EMG activity. Analysis showed no main effect of gaze condition for triceps brachii for both Early and Late Perturbations [Early: F(1,31) = 3.2, p = 0.66, η² = 0.5; Late: F(1,31) = 0.7, p = 0.9, η² = 0.01], left tibialis anterior [Early: F(1,31) = 4.1, p = 0.9, η² = 0.49; Late: F(1,31) = 1.1, p = 0.53, η² = 0.04], or right tibialis anterior [Early: F(1,31) = 2.8, p = 0.2, η² = 0.5; Late: F(1,31) = 0.8, p = 0.76, η² = 0.01]. Figure S2 shows comparable EMG activity between pursuit (SPEM) and fixation (FC) conditions for the three muscles.

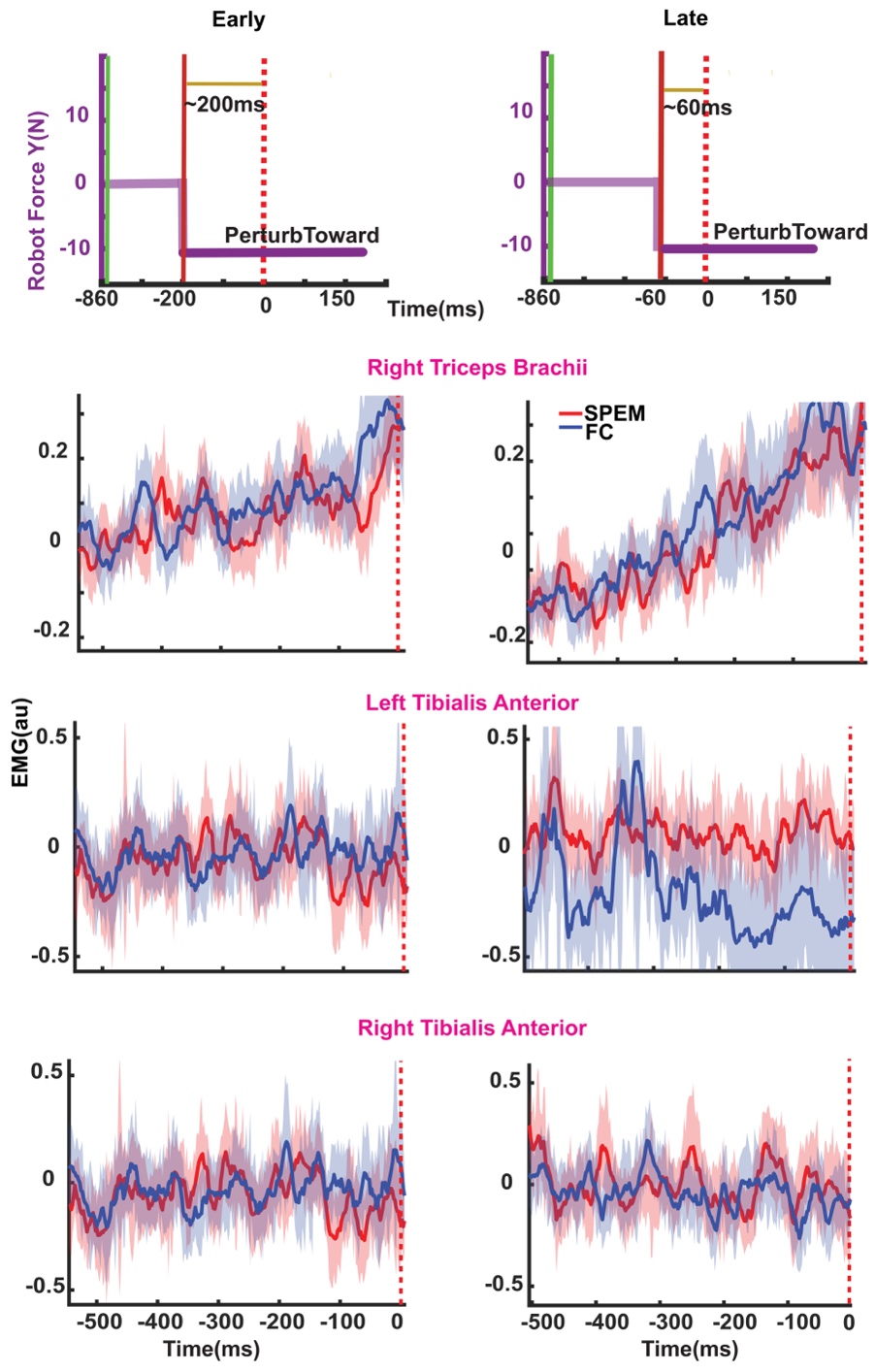

Figure S2: Pre-perturbation EMG Activity. EMG recordings during early (Early, ~200 ms pre-contact) and late (Late, ~60 ms pre-contact) perturbation trials. (A) Right triceps brachii, (B) left tibialis anterior, and (C) right tibialis anterior show comparable anticipatory activity between pursuit (red) and fixation (blue) conditions.

**Center of pressure excursions were similar across the gaze conditions**

The center of pressure excursions along the Y direction (direction in which the perturbations were applied) were similar between the two gaze conditions at both the time points. Figure S3 shows the center-of-pressure trajectory during during early (Early, ~200 ms pre-contact, left panel) and late (Late, ~60 ms pre-contact, right panel) perturbation trials.

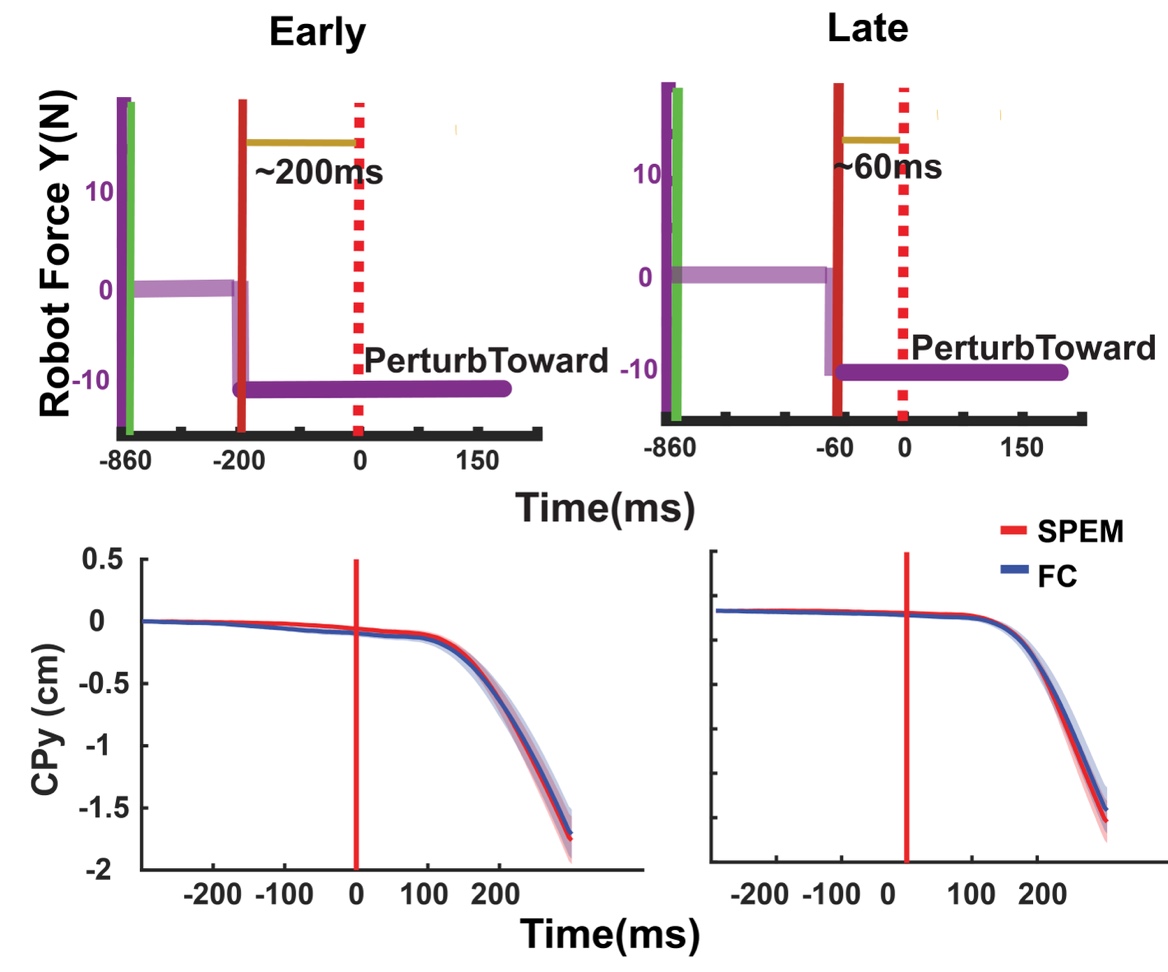

Figure S3: Center of pressure (Y-axis) shows no gaze-dependent differences at either perturbation timing.
